# In planta transposon sequencing for virulence gene identification in *Xylella fastidiosa*

**DOI:** 10.1101/2024.09.23.614259

**Authors:** Lindsey Burbank, Elizabeth Deyett, Nancy Her, Sydney Helm Rodriguez, Mayra Magdeleno, Philippe E. Rolshausen, Caroline Roper

## Abstract

In bacterial genetics, large-scale screening approaches such as sequencing transposon mutant pools can be highly effective for identifying and characterizing genes with unknown functions. In the plant pathogen, *Xylella fastidiosa*, this approach is challenging due to the fastidious nature of this bacterial species and its niche-specific growth in the plant xylem tissue. The purpose of this study was to explore the feasibility of transposon sequencing (Tnseq) for identification of virulence genes in *X. fastidiosa*, with the hypothesis that this would uncover genes or pathways not previously associated with plant infection. Predicted essential genes were compared after *X. fastidiosa* strain M23 was grown *in vitro* and *in planta* using two known susceptible host species (grapevine and almond). After growth *in planta*, several gene categories were predicted as essential including hemagglutinins, tRNAs, toxin-antitoxin systems, and prophage genes. Three predicted essential genes (XfasM23_0359, XfasM23_0360, XfasM23_0972) were chosen for further validation by making targeted deletion mutants. Deletion mutants exhibited reduced disease in grapevines, but normal growth and aggregation phenotypes *in vitro*. Overall, the Tnseq approach has some practical limitations due to the nature of the *X. fastidiosa* pathosystem, and significant bottleneck effects of inoculation, but was still able to identify genes contributing to disease in plants. Recommendations for future Tnseq studies in *X. fastidiosa* are discussed based on the challenges and results of this work.

**Importance:** *Xylella fastidiosa* is a plant pathogenic bacterial species that causes significant economic damage in multiple agricultural industries. Globally, disease epidemics in citrus, grapes, almonds, and olives launched widespread efforts in pathogen surveillance, and quarantine restrictions on plant commodities. Research efforts on *X. fastidiosa* biology and pathogenesis have still not yielded many new and effective disease control measures, and management in most areas relies primarily on insect vector control. Expansion of available genetic research tools to include high-throughput mutant screening protocols for *in planta* experiments will facilitate identification of novel disease control targets for this pathogen.

## Introduction

*Xylella fastidiosa* causes disease in multiple divergent plant species, several of which are important agricultural commodities. Currently found in the Americas, and parts of Europe and Asia, *X. fastidiosa* is a significant issue for plant production and trade (1). Major crops impacted include grapes, citrus, almond, and olive among others. *X. fastidiosa* is transmitted between plants by several different insect species, including sharpshooter leafhoppers, and spittlebugs (2, 3). Invasion into new areas can occur by movement of infected plant material or insect vectors, and may go undetected for months or years due to slow disease progression and symptoms that mimic drought or other stress conditions (4). This makes disease control challenging since the initial identification of an outbreak may reveal prior undetected pathogen spread. Development of resistant plant material and control of insect vectors are currently the most effective strategies for managing *X. fastidiosa* diseases in agriculture, but neither of these approaches are sufficient on their own (4). Environmental concerns and increasing insecticide resistance place limitations on large scale vector control (5, 6), and plant breeding efforts require significant time and resources (7). To make disease management at the plant level more adaptable, scaling up identification of pathogen processes that can be targeted for preventing infection is important.

One of the intriguing aspects of *X. fastidiosa* infection is the ability of this pathogen to reside asymptomatically in a wide range of plant species with only specific combinations of pathogen strains and plant hosts resulting in severe disease outcomes (8). With greater availability of bacterial genome sequences in recent years, more extensive analysis is possible to predict genes important for evolution, host adaptation, and pathogenic interactions (9, 10). Large scale functional screening such as using random transposon libraries complements genomic approaches in accelerating insights into pathogen interactions with the infected host. The benefit of high-throughput screening applications in bacterial-host interactions is in identifying novel pathways and functions that would not have been predicted based on prior knowledge. Transposon sequencing (Tnseq) protocols have been used in several plant pathogenic bacteria to uncover previously unknown infection-mediating pathways. In *Dickeya dadantii*, a Tnseq strategy established the importance of biosynthesis pathways for purine nucleotides, as well as amino acids leucine, cysteine, and lysine in colonization of macerated plant tissues (11). In *Pseudomonas syringae*, this approach was also used to compare fitness-associated genes in three different host plants (common bean, lima bean, and pepper), identifying alginate polysaccharide biosynthesis and phytotoxin production as virulence traits that differ across hosts (12). *X. fastidiosa* specifically infects the xylem vessels of plants and is naturally transmitted via sap-feeding insects. This mode of transmission necessarily creates bottlenecks in the bacterial population as only a small number of cells are likely to be transferred in a single inoculation event. Experimentally, xylem puncture methods are typically used for *X. fastidiosa* inoculation to deposit the bacteria directly into the plant vascular system. Although mechanical xylem puncture can deposit more bacterial cells directly into the xylem, in is not clear how many of those cells successfully gain entry and propagate throughout the xylem vessels. In *Pantoea stewartii*, another xylem-infecting pathogen, Tnseq experiments were conducted using similar inoculation methods and found that ∼10% of the inoculated cells became established in the plant (13). Multiple pathways associated with fitness in the xylem for *P. stewartii* were still identified including iron uptake, outer membrane proteins, and transport pathways (13). Similarly, cell envelope functions and *de novo* amino acid synthesis contributed significantly to fitness of *Ralstonia solanacearum* in tomato xylem sap compared with rich growth medium in Tnseq comparisons (14). Tnseq would be a useful tool to advance understanding of *X. fastidiosa*-host interactions and disease development dynamics if it can be implemented effectively. The goal of this study was to explore the use of Tnseq in *X. fastidiosa* for *in planta* study of virulence and fitness associated genes, and to provide adapted protocols and information on this approach to facilitate further research. With some limitations, mainly due to significant bottlenecks, Tnseq identified several previously uncharacterized genes that may impact *X. fastidiosa* fitness during plant infection. Furthermore, three of these genes were experimentally confirmed to encode functions involved in virulence. Recommendations for *X. fastidiosa* Tnseq studies are also discussed for consideration in future research.

## Materials and Methods

### Tn5 library construction

*X. fastidiosa* subsp. *fastidiosa* strain M23, originally isolated from almond trees showing leaf scorch disease in Kern County, CA (15) was used as a background for Tn5 mutagenesis. Wild type M23 was transformed by electroporation with 1 µl of EZ Tn5 <KAN-2>Tnp Transposome (LCG Biosearch Technologies) as previously described (16). One µl of TypeOne Restriction Inhibitor (LCG Biosearch Technologies) was added to the transformation reactions. Three separate transformation reactions were performed, and each was recovered in 1 ml PD3 for 16 hours on a shaker at 180 rpm. After the recovery period, 20 µl of each transformation was removed and used for serial dilution plating to determine the number of transformants. Entire transformation volumes from all three reactions were plated on PD3 plates with kanamycin (5 µg/ml) and incubated at 28°C for 10 days. Based on concentration of transformants determined by dilution plating and total volume of transformation reactions, ∼10,000 transformants were obtained for each transformation. All transformants were pooled, and the combined mutant library was saved in 15% glycerol stocks at −80°C.

### Plant inoculation of pooled transposon mutants

Wild type M23 and pooled mutant library were grown on PD3 plates (with kanamycin for mutants) for 6 days at 28°C. Cells were harvested by scraping entire plate with 1xPBS. Five plates were combined for each strain, and cell concentration was normalized to OD_600nm_ = 0.5. A total of 12 ml of cell suspension was obtained for each strain. Cell suspension was centrifuged at 5000 rpm for 5 minutes to pellet the cells and buffer was removed. Cells were resuspended in 0.1 volume (1.2 ml) of 1xPBS to create a highly concentrated cell suspension (OD_600nm_∼5.0). This cell concentration is significantly higher than is generally used for *X. fastidiosa* inoculation, but was used to increase the number of mutants introduced to into the plant. Potted two-year-old grapevines (*Vitis vinifera* cv Chardonnay) and almond seedlings (Prunus *dulcis* Y113-20) were inoculated by pinprick with 100 µl total of cell suspension (four drops of 25 µl). Four plants of each species were inoculated with each strain (wild type M23 and transposon mutant library) and four plants of each species were inoculated with 1xPBS to serve as negative controls. Plants were inoculated in mid-summer (early July 2019).

### Estimation of bottleneck effect

Grapevines were inoculated with wild type *X. fastidiosa* M23 using the same method described above for introduction of mutant libraries into plants. In addition to the high concentration of cells used for inoculation of Tn5 libraries (OD_600nm_∼5.0), separate plants were inoculated with a cell concentration of OD_600nm_∼0.25 which is the typical cell concentration used to test virulence in *X. fastidiosa*. Serial dilutions of the two different cell concentrations were plated on PD3 plates to quantify colony forming units (CFU) in the inoculum. Three plants were inoculated with each cell concentration and a total of 80 µl of inoculum was introduced into each plant. After 24 hours, a 5 cm stem section surrounding the point of inoculation was cut and ground up in 2 ml 1xPBS. Serial dilutions were plated on PD3 plates and incubated at 28°C for one week. CFUs from the initial inoculum and from the plant samples were compared for an estimate of the percentage of cells established in the plant (Supplemental Table S1). The bottleneck estimated that <1% of the inoculated cells were established in the plant.

### Bacterial isolation and DNA extraction

After ten weeks post-inoculation, bacterial populations were isolated from each inoculated plant. For grapevines, petioles from leaves with scorching symptoms were selected from lower, middle, and upper sections of each plant and combined. For almond, leaves with scorching symptoms were selected from lower, middle, and upper sections of each plant and midrib was excised with a sterile razor blade. Only symptomatic leaves were sampled to maximize the bacterial population obtained from the plant. All samples were surface sterilized in 70% ethanol for 30 seconds, 20% bleach for 30 seconds, and washed twice in sterile water. After sterilization, all samples from the same plant were combined in a mesh grinding bag (Agdia Inc) and crushed with two ml of 1xPBS. Plant extract was collected, and five serial dilutions were made in 1xPBS. Twenty µl of each dilution was plated separately to quantify bacterial populations and the rest of the total volume was plated on PD3 plates and grown for 10 days. Cells were harvested by scraping off plates in 1xPBS. Bacterial cells from the same plant were all combined and cell concentration was normalized to OD_600nm_ = 0.5. One ml aliquots were centrifuged at 9000 rpm for 5 minutes, supernatant was removed, and cell pellets were frozen at −20°C until DNA extraction. DNA was extracted using a DNeasy Blood and Tissue Kit (Qiagen) with protocol as follows, slightly modified from the manufacturer’s instructions.

Frozen cell pellets were resuspended in 180 µl buffer ATL with 20 µl proteinase K added. Samples were incubated at 56°C for 1 hour, mixing every 15 minutes by vortex. After incubation, 20 µl of 20 mg/µl RNase A (ThermoFisher) was added to each sample and incubated at room temperature for 15 minutes. Buffer AL (200 µl) and 100% ethanol (200 µl) were then added to each sample and the entire volume was transferred to spin columns. After washing twice with the kit provided wash buffers, samples were eluted with diH_2_O heated to 37°C (50 µl x 2). DNA samples from the same plant were combined and concentrated with a Speedvac concentrator to a final concentration of 100-500 ng/µl. From the four plants each of almond and grapevine, the three plants from each species which yielded the highest amount of combined bacterial DNA was moved forward for sequencing. Three samples of the original pool of transposon mutants that was grown on PD3 plates for one weeks were also sequenced as control samples.

### Sequencing and QC

Custom library construction and Illumina sequencing targeting the Tn5 transposon insertion regions was performed by Fasteris SA (Geneva, Switzerland) using individual indexing. Total of 3 µg DNA for each of the nine samples (3 grapevine, 3 almond, 3 PD3 medium) was provided for this process and DNA quantity and quality was confirmed by gel electrophoresis and fluorescent quantification using Quant-IT DNA quantification kit (ThermoFisher). Although the library preparation pipeline is proprietary, similar sequencing protocols are available elsewhere (17). Sequencing of library pool was run on NovaSeq 6000, SP-100, with customized run of 1×75 bp single reads. A 10% PhiX internal control was included in the sequencing run. Sample indexes and raw sequence quality metrics are included in supplemental Table S2.

### Transposon insertion site mapping and essential gene analysis

Raw sequence reads were filtered by transposon sequence (supplemental figure S1) and trimmed for adaptor removal with Cutadapt (18). Reads were then mapped to the *X. fastidiosa* M23 reference genome (NCBI Accession CP001011.1) using STAR aligner (19). Identification of transposon insertion sites and essential gene prediction was performed with the Bio-TraDIS pipeline (20). Computational prediction of essential genes from Tnseq data is based on the frequency of specific mutants in the pool of sequenced insertion sites. Using the Bio-TRADIS analysis pipeline, sequence reads containing insertion sites were mapped to the reference genome to identify the number of unique insertion sites per sample, which was then normalized to gene length to give an insertion index (20). Based the frequency distribution of the insertion index for each gene, likelihood ratios were calculated for whether genes are more likely to be essential (no or low frequency of insertions present) or non-essential (high frequency of insertions present). The defined cutoff for essentiality used by the Bio-TRADIS pipeline is a log2-likelihood ratio of less than −2 (a gene is at least four times more likely to be essential than not)(20, 21). The insertion index corresponding to this cutoff was calculated for each sample (Insertion index for Essentiality, Table 1), and genes with an insertion index below that are referred to as “predicted essential genes” as defined by the Bio-TRADIS pipeline (Table 1). Supplemental Figure S2 shows PcOA plots of predicted essential genes by treatment (control vs grapevine vs almond).

**Table 1.** Read mapping and transposon insertion sites.

| Sample* | Mapped Reads | % Uniquely mapped reads | % Mapped reads | Sequenced bases | No. of Unique Insertion Sites | Insertion Index for Essentiality | Number of Predicted Essential Genes | Average Transposon / X bp |
| --- | --- | --- | --- | --- | --- | --- | --- | --- |
| PD3-1 | 27392901 | 93.8% | 98.4% | 1190324111 | 51622 | 0.0011 | 142 | 49 |
| PD3-2 | 38867977 | 94.9% | 98.2% | 1720191336 | 66950 | 0.0043 | 135 | 38 |
| PD3-3 | 32846188 | 94.4% | 98.3% | 1450784298 | 53619 | 0.0018 | 123 | 47 |
| GR-1 | 31284962 | 93.5% | 98.3% | 1370535739 | 78289 | 0.0075 | 129 | 32 |
| GR-2 | 30526989 | 96.5% | 99.0% | 1373669949 | 37036 | 0.0003 | 175 | 68 |
| GR-3 | 38870221 | 94.3% | 98.5% | 1728972909 | 58952 | 0.0044 | 142 | 43 |
| AL-1 | 24719460 | 69.5% | 72.1% | 1101268541 | 47442 | 0.0020 | 163 | 53 |
| AL-2 | 35441252 | 96.1% | 98.2% | 1585732291 | 39595 | 0.0010 | 165 | 64 |
| AL-3 | 45922319 | 96.9% | 98.4% | 2059390753 | 41080 | 0.0010 | 162 | 62 |
| **Totals/Averages** | 33985808 | 92.2% | 95.48% | 13580869927 | 52732 | 2.70E-03 | 148 | 51 |

\*PD3 = cells grown in culture, GR = grapevine, AL = almond

### Site-directed mutagenesis and complementation

Gene deletion constructs were created by amplifying the adjacent 5’ and 3’ flanking regions and removing the gene of interest with a replacement of a kanamycin resistance cassette using overlap extension polymerase chain reaction (PCR). All flanking regions and kanamycin resistance cassettes were amplified using the manufacturer’s protocol for Q5® High-Fidelity DNA Polymerase (New England Biolabs) and with primer concentrations of 10μM. The 5’ flanking region for M23-0359 at the size of 916bp was amplified using the primer pairs 5GPWF/5GPW RKAN at a melting temperature (Tm) of 68°C. For the 5’ flanking region of M23-0360 (at 575bp) was amplified using the primer pairs 5PMSKF/ 5PMSKRKAN at Tm of 63°C. The primer pair 5M23-0972F/5M23-0972RKAN was used to amplify the 5’ flanking region of M23-0972 (792bp) at Tm of 57°C. The 3’ flanking region for M23-0359 (751bp) was amplified using GPWR/3GPWFKAN at Tm of 68°C, for M23-0360 (986bp) using primer pair 3PMSKR/3PMSKFKAN at Tm of 63°C, and for M23-0972 (871bp) using the primer pair 3M23-0972R/3M23-0972FKAN at Tm of 54°C. The kanamycin resistance cassette was amplified from the pCR™-Blunt vector (Invitrogen) using primer pair KANF5GPW/KANR3GPW for XfasM23_0359 at Tm of 61°C, KANF5PMSK/KANR3PMSK for XfasM23_0360 at Tm of 59°C, and KANF5M23-0972/KANR3M23-0972 for XfasM23_0972 at Tm of 61°C. The respective kanamycin resistance cassettes replaced the open reading frame of the XfasM23_0359 gene from nucleotide 456911 to nucleotide 457249, XfasM23_0360 gene from nucleotide 457447 to nucleotide 457728, and XfasM23_0972 gene from nucleotide 1135138 to nucleotide 1135449. The three fragments for each construct were annealed at equimolar amounts at 98°C for 30s entering a 10 cycle rotation of 98°C for 10s, at 69°C for 30 secs, and 72°C for 1 min. Immediately after, 1.25μL of the flanking primer pairs 5GPWF/3GPWR for M23-0359, 5PMSKF/3PMSKR for M23-0360, and 5M23-0972F/3M23-0972R for M23-0972 were added including 0.25μL of Q5® High-Fidelity DNA Polymerase (New England Biolabs) to their respective reactions and amplified under these conditions: 98°C for 30s and following 30 cycles of 98°C for 10s, 25s at 68°C for M23-0359 and M23-0369 and 57°C for M23-0972, and 72°C for 25s with a single final extension for 20min at 72°C. The resulting amplicons (M23-0359 at 2.559kb, M23-0360 at 2.457kb, and M23-0972 at 2.559kb) were ligated into the cloning vector pCR8/GW/TOPO (ThermoFisher) using the manufacturers protocol to create the Δ*0359* deletion construct, Δ*0360* deletion construct, and Δ*0972* deletion construct. Plasmid constructs were transformed into *X. fastidiosa* M23 by electroporation (16) and transformants selected on PD3 plates with kanamycin. Correct mutations were confirmed by PCR amplification of the mutated region (primers M23-0359-F/M23-0359-R, M23-0360-F/M23-0360-R, M23-0972-F/M23-0972-R) and Sanger sequencing of the PCR product.

For gene complementation, open reading frame of the gene of interest plus 250bp upstream to include the promoter region was PCR amplified from wild type M23 DNA (primer pairs: 0359-ORF-F/0359-ORF-R, 0360-ORF-F/0360-ORF-R, 0972-ORF-F/0972-ORF-R). Because XfasM23_0360 is a putative toxin gene as part of a toxin-antitoxin system, complementation construct for this mutant included both the toxin and antitoxin (XfasM23_0360 and XfasM23_0361) to avoid toxicity of the plasmid constructs. PCR products were transformed into cloning vector pCR8/GW/TOPO and confirmed by Sanger sequencing. Then each complementation construct was recombined with plasmid pAX1/GW to create *X. fastidiosa* genomic insertion constructs using LR recombinase (Invitrogen). These constructs of pAX1 were transformed into respective deletion mutants by electroporation for genomic insertion of the wild type gene copy. Transformants of complemented strains were selected on gentamycin. All plasmids and bacterial strains are listed in supplemental Table S3. All primers sequences used for mutagenesis and complementation are listed in supplemental Table S4.

### Grapevine inoculations with deletion mutants

Wild type *X. fastidiosa* M23, deletion mutants, and complemented strains were inoculated into potted two-year-old grapevines (*Vitis vinifera* cv Chardonnay) using the same procedure as described for the transposon mutant pool except with a lower inoculum dose. For inoculation of individual mutants, cells were suspended at a concentration of OD_600nm_=0.25 (∼10^8^ cfu/ml) and each plant was inoculated with 80 µl of cell suspension. Fifteen plants were inoculated per bacterial strain and 15 uninoculated grapevines were included as negative controls. Plants were randomized within the greenhouse and maintained at temperatures between 20°C and 37°C. Disease progression was evaluated using a disease scoring system as previously described where 0 = no PD symptoms, 1 = one or two leaves just beginning to show marginal necrosis, 2 = two to three leaves with significant marginal necrosis, 3 = one half or more of the leaves showing marginal necrosis and a few match sticks, 4 = all of the leaves showing heavy scorching and numerous matchsticks, and 5 = a dead vine (22). Disease ratings were done blind to the treatments to reduce rater bias. Petiole samples were collected 8 weeks after inoculation for confirmation of infection and quantification of bacterial populations. Three petioles were sampled from each plant, one each from the bottom, middle, and upper thirds of the vine. Petioles from the same plant were pooled together as one sample for further processing. Samples were lyophilized using a FreeZone freeze dryer (LabConco) and pulverized using a TissueLyzer II (Qiagen) with tungsten beads. DNA was extracted from pulverized tissue using a CTAB/phenol/chloroform extraction and isopropanol precipitation as previously described (23). DNA was resuspended in TE buffer and diluted 1:10 in sterile distilled water for qPCR quantification. Stem samples were collected at the termination of each experiment by cutting a 2.5 cm section of the grapevine main stem from 30 cm above the point of inoculation. Stem samples were frozen at −20°C until DNA extraction which was carried out using the same protocol as for petiole samples. Quantification was performed using Applied Biosystems Fast SYBR Green Master Mix (ThermoFisher) and primers Xf145-60F/Xf145-60R based on a standard curve created from serial dilutions of *X. fastidiosa* DNA (known concentration in cfu/ml) combined with plant DNA (24). Bacterial quantity was normalized to total DNA concentration obtained from fluorescent quantification using Quant-IT DNA quantification kit (ThermoFisher).

### *In vitro* growth and cell aggregation measurements

All strains were grown on PD3 plates for 6 days. Cells were scraped off plates in liquid PD3 medium and adjusted to a concentration of OD_600nm_=0.25. Then 500 µl of cell suspension was added to 5 ml PD3 in 15 ml polypropylene test tubes and incubated statically at 28°C for 14 days. After 14 days, OD_600nm_ was measured of the undisturbed cultures (OD), cultures were mixed thoroughly by vortexing and pipetting, and OD_600nm_ was measured of the total culture (ODt) after mixing. Percent aggregation was calculated as: [(ODt-OD)/ODt]x100 as previously described (23). Each strain was measured twice per experiment and the experiment was run independently three times. To evaluate total growth rate, the OD_600nm_ of the cultures after mixing (ODt) was compared at 7 and 14 days of growth.

### Toxin-antitoxin gene cloning

Open reading frames of the toxin (primers: 0360-ORF-F/0360-T-rev), antitoxin (primers: 0361-AT-fwd/0361-AT-rev), and both together (primers: 0360-ORF-F/0361-AT-rev) were PCR amplified using a polymerase that creates blunt-ended products (Phire Plant Direct PCR master mix, ThermoFisher). PCR products were evaluated for correct size by gel electrophoresis and 1 ul was cloned into a blunt-cloning vector that has counter selection to prevent empty vector ligation (Zero Blunt PCR cloning kit, ThermoFisher). Negative control cloning reaction was included that contained no DNA insert. Serial dilutions were plated on LB with kanamycin and incubated at 37°C overnight. Colonies were counted, and three clones for each construct were confirmed by Sanger sequencing. Five replicates of the cloning were performed.

### Statistical Analysis

All statistical analyses and graphical representations were done using OriginPro2023 software package (OriginLabs Inc). Quantitative datasets were evaluated for normal distribution using Shapiro-Wilkes test at 0.05 significance level. Datasets that follow a normal distribution were evaluated using one-way ANOVA and post-hoc Tukey means comparison tests at the 0.05 significance level. Datasets that did not follow normal distribution were evaluated with non-parametric Kruskal-Wallis ANOVA and Dunns test. Details of statistical analysis are included in Supplemental File 2.

## Results

### Prediction of essential genes in *X. fastidiosa in planta*

Analysis of sequence reads from random transposon mutant pools of *X. fastidiosa* grown *in vitro* (PD3 medium) or *in planta* (grapevine and almond) predicted an average of 148 essential genes as defined by the Bio-TRADIS output terminology across the different conditions (Table 1). Because these predictions likely over-estimate essentiality due to bottlenecks caused by inoculation and variability between samples (i.e. mutants not present due to chance), the list of predicted essential genes in a specific condition was filtered to include genes predicted to be essential in all three replicates of that condition but not in the other conditions (Fig. 1, Table S5). In total, 9 genes were predicted as “essential” in PD3 growth condition but not in either plant condition, 16 genes were predicted as essential only in grapevine, and 27 genes only in almond (Fig. 1). This comparison was performed to evaluate gene functions that might be important in a specific condition. Of the predicted “essential” protein-coding genes (excluding rRNA and tRNA genes), the majority were annotated as hypothetical proteins. However, several other gene categories were also identified including hemagglutinins, phage-associated genes, and conjugal transfer proteins (Fig. 1). In general, the almond growth condition had less variability between samples compared with the grape treatment, so more genes were found in all three replicates in almond. Separately, predicted essential genes *in planta* were compared with *in vitro* using predictions from almond and grapevine combined. Genes listed in Table 2 were predicted as “essential” in at least five out of the six plant samples but not in the PD3 samples.

**Figure 1.**
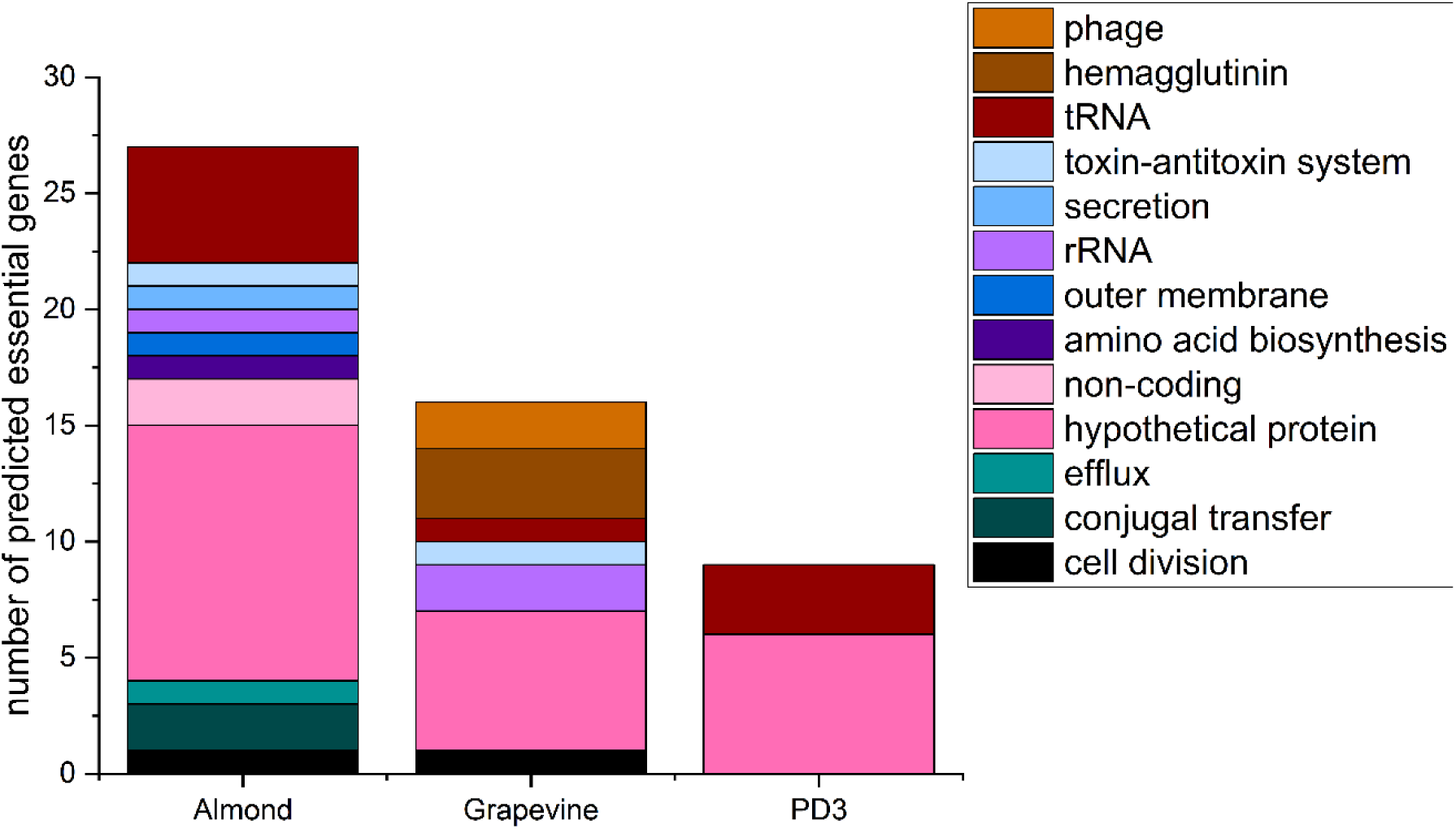
Predicted essential genes under different growth conditions. Number of predicted essential genes after bacterial propagation in grapevine, almond, or culture medium (PD3). Genes listed are predicted as “essential” in all replicates of the specified treatment but not under the other conditions. Genes were categorized based on functional predictions from the reference genome annotation.

**Table 2.**
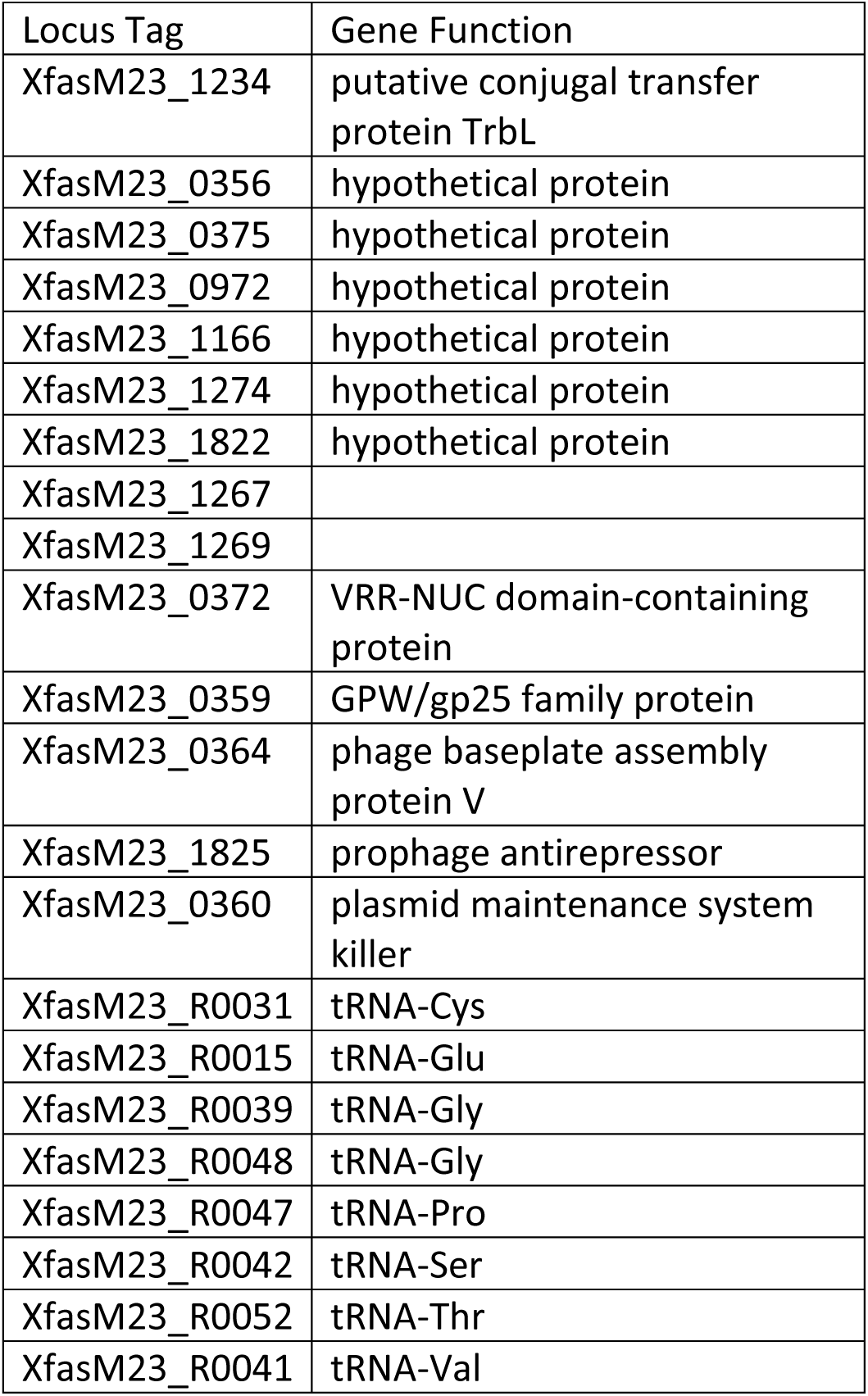
Predicted essential genes *in planta*.

| Locus Tag | Gene Function |
| --- | --- |
| XfasM23_1234 | putative conjugal transfer protein TrbL |
| XfasM23_0356 | hypothetical protein |
| XfasM23_0375 | hypothetical protein |
| XfasM23_0972 | hypothetical protein |
| XfasM23_1166 | hypothetical protein |
| XfasM23_1274 | hypothetical protein |
| XfasM23_1822 | hypothetical protein |
| XfasM23_1267 |  |
| XfasM23_1269 |  |
| XfasM23_0372 | VRR-NUC domain-containing protein |
| XfasM23_0359 | GPW/gp25 family protein |
| XfasM23_0364 | phage baseplate assembly protein V |
| XfasM23_1825 | prophage antirepressor |
| XfasM23_0360 | plasmid maintenance system killer |
| XfasM23_R0031 | tRNA-Cys |
| XfasM23_R0015 | tRNA-Glu |
| XfasM23_R0039 | tRNA-Gly |
| XfasM23_R0048 | tRNA-Gly |
| XfasM23_R0047 | tRNA-Pro |
| XfasM23_R0042 | tRNA-Ser |
| XfasM23_R0052 | tRNA-Thr |
| XfasM23_R0041 | tRNA-Val |

### Evaluation of virulence role for select predicted essential genes

Three predicted essentialgenes were selected for further testing: XfasM23_0972 (hypothetical protein), XfasM23_0359 (phage associated protein), and XfasM23_0360 (toxin component of toxin-antitoxin system). These particular genes were selected because they were predicted “essential” in all at least five of the six plant samples sequenced, and they did not have previously described roles in *X. fastidiosa* virulence. Targeted deletion mutants (Δ*0359*, Δ*0360*, Δ*0972*) for these three genes caused decreased leaf scorching symptoms in grapevine compared to the M23 parent strain when inoculated individually (Fig. 2A). Complemented mutants included in a second replicate of virulence testing caused similar levels of disease symptoms compared with wild type M23 (Fig. 2B).

**Figure 2.**
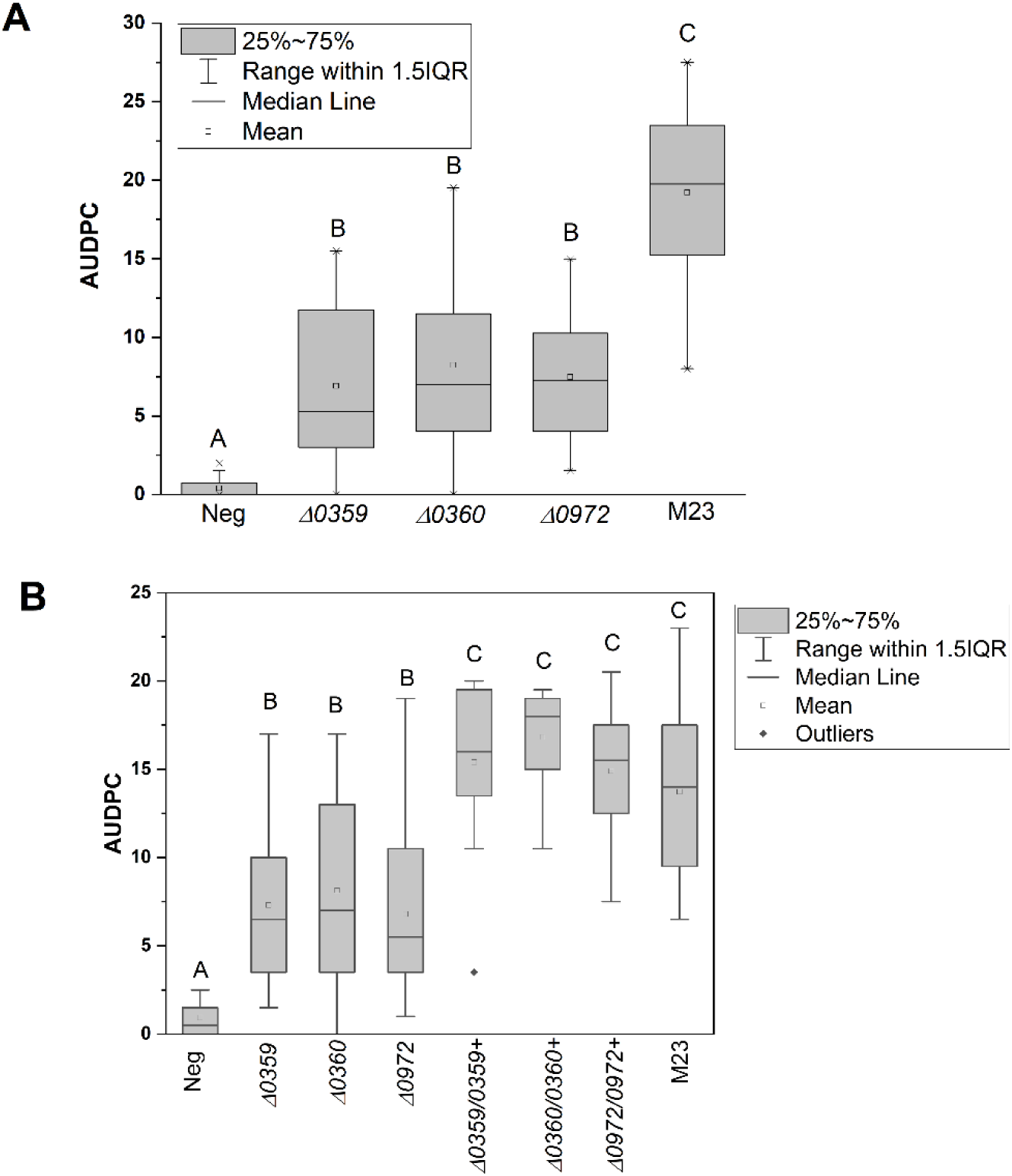
Disease symptoms of targeted deletion mutants. A) Deletion mutants (Δ0359, Δ0360, Δ0972) and wild type M23 were inoculated into susceptible grapevines (*Vitis vinifera* Chardonnay) by pinprick inoculation and evaluated weekly for scorching symptoms using a 0-5 disease scoring system. Disease scores over time were used to calculate area under the disease progress curve (AUDPC) using the agricolae package in R. Different letters above boxes indicate significant difference based on ANOVA and post-hoc Tukey means comparison testing at the 0.05 significance level. B) Inoculation experiments were repeated a second time following the same process, but with the inclusion of complemented strains for each mutant.

### Cell aggregation and growth rate of deletion mutants

Deletion mutants Δ*0359*, Δ*0360*, and Δ*0972* were evaluated for growth abnormality *in vitro*. The characteristic growth pattern of *X. fastidiosa* is cell aggregation in liquid cultures. All three of the deletion mutants and respective complemented strains showed the same growth habit as the wild type M23 strain (Fig. 3A).

**Figure 3.**
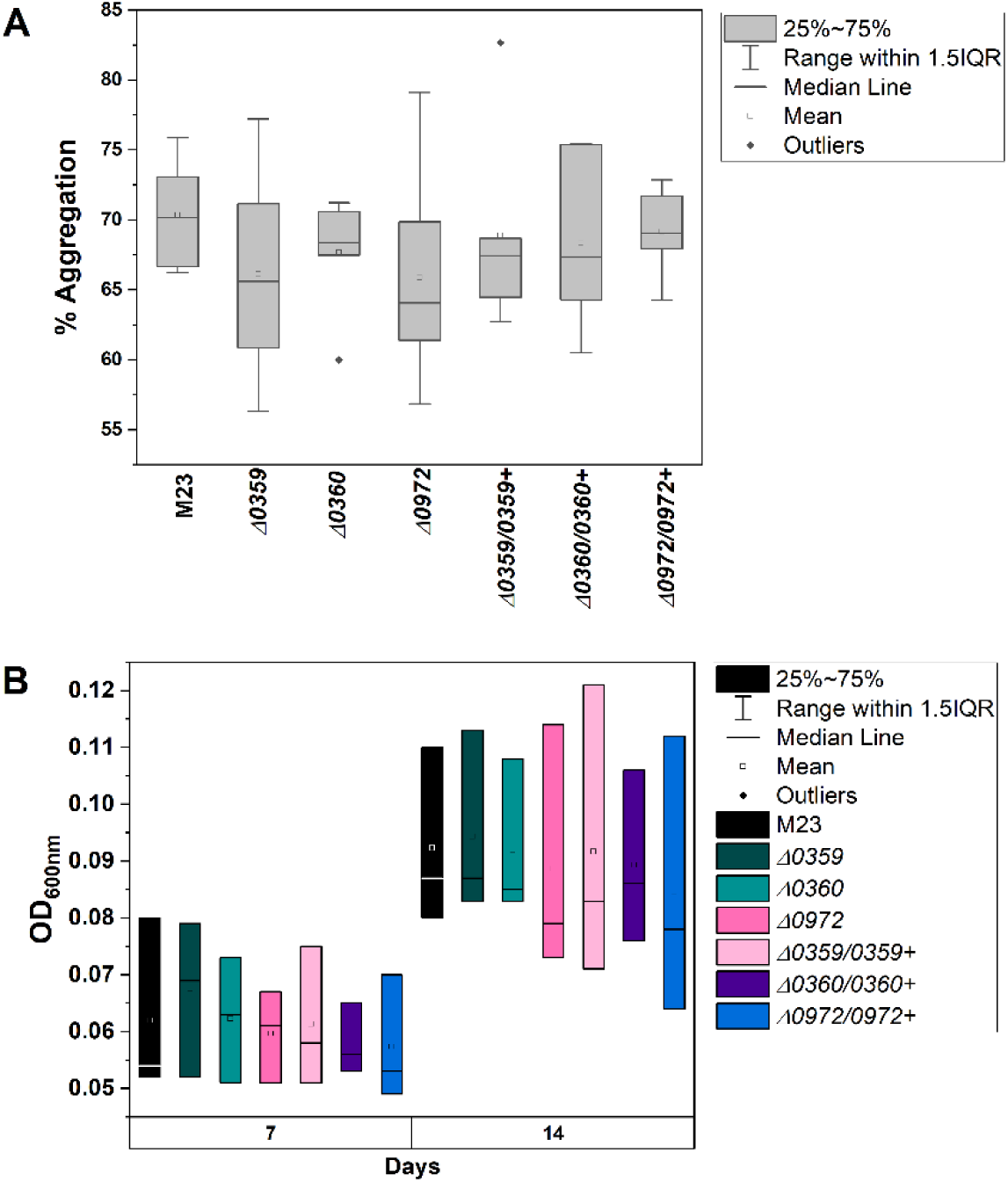
Cell aggregation and in vitro growth of deletion mutants. Deletion mutants (Δ*0359*, Δ*0360*, Δ*0972*) and respective complemented strains were grown in liquid PD3 medium and evaluated for growth rate and cell aggregation over 14 days. A) Cell aggregation percentage is calculated based on OD_600nm_ of dispersed and total cells (OD_600nm_ after thorough mixing) as previously described (24). B) Cell growth is indicated by OD_600nm_ of total cells after 7 and 14 days of growth at 28°C. For both aggregation and cell growth, duplicate measurements were taken of each culture, and experiments were repeated independently three times.

Growth rate in rich medium (PD3) was also not impacted in the deletion mutants over the course of two weeks in culture (Fig. 3B). Bacterial quantity was also determined in inoculated plants to assess whether growth of the bacterial population was altered in the deletion mutants. Petiole and stem samples from inoculated grapevines were tested with qPCR after eight weeks post-inoculation (Fig. 4). There was no significant difference in average bacterial population in mutant-inoculated plants compared with the wild type M23 strain based on petiole samples (Fig. 4A) or stem samples (Fig. 4B). However, there was greater sample-sample variability in petiole samples from mutant inoculated plants than for the wild type or complemented strains (Fig. 4A). A similar trend was observed in sample-sample variability in bacterial quantity detected in almond petiole samples (Supplementary Fig. S3). Stem samples were less variable overall, and did not show reduced growth of the mutants in plants (Fig. 4B). More in depth information on the biological functions and additional characterization of the three genes tested is included in Supplemental File 3.

**Figure 4.**
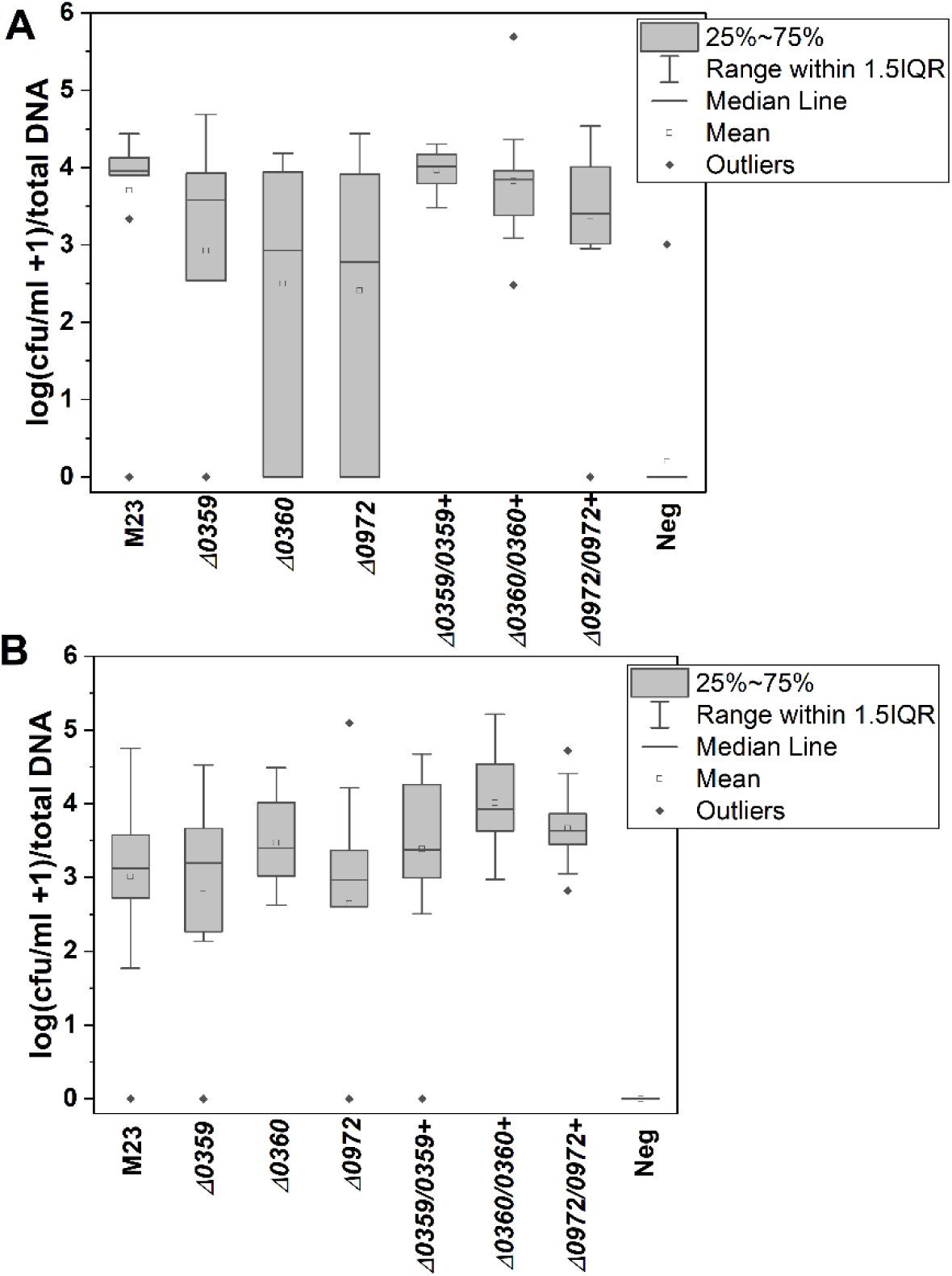
*Xylella fastidiosa* growth in plants. Quantity of *X. fastidiosa* was evaluated by qPCR after inoculation of wild type M23, deletion mutants, and respective complemented strains in susceptible grapevines. A) *X. fastidiosa* quantity in petiole samples after 8 weeks post-inoculation B) *X. fastidiosa* quantity in stem sections taken from 30 cm above the point of inoculation after 12 weeks post-inoculation.

## Discussion

The infection process for *X. fastidiosa* in plants is initiated by inoculation into the xylem vessels by sap-feeding insects (2, 3). Once within the xylem, *X. fastidiosa* can move throughout the plant via twitching motility (25) and cross between xylem vessel segments through the production and secretion of cell wall degrading enzymes (22, 26, 27). During this process, *X. fastidiosa* will attach and form biofilms in xylem vessels (28, 29), and must evade recognition by the plant immune system (30–32), as well as maintaining necessary metabolic functions (33). To be transmitted to a new host plant, *X. fastidiosa* can be picked up by sap-feeding insects, adhering to the inside of the insect mouthparts as they consume xylem sap (34). Genetic components necessary for all these processes in *X. fastidiosa* infection have been characterized mainly by mutagenesis experiments and phenotyping of individual deletion mutants. However, this approach relies on accurately predicting genes likely to be important for plant infection, and is time and resource intensive due to the fastidious nature of *X. fastidiosa* and the fact that many of its host plants are woody perennials. Tnseq has the potential to broaden the scope of fundamental research on *X. fastidiosa* and identify previously unknown aspects of the pathogenesis process, but comes with some limitations and challenges in this particular pathosystem. Bottleneck effects during inoculation and re-isolation of the bacterial population are always a factor in Tnseq experiments (35). This is especially the case with *X. fastidiosa* in plants because it needs to be inoculated directly into the xylem vessels by needle puncture which limits the volume of bacterial suspension that can be used for inoculation. Using standard needle-inoculation methods, based on re-isolating from the point of inoculation after 24 hours, <1% of *X. fastidiosa* cfu established in grapevine stems causing a significant bottleneck. Increasing the inoculum concentration from ∼10^8^cfu/ml to ∼10^12^cfu/ml did increase the number of established cells, but only by about 100-fold. It is likely that the true number of established cells is higher than estimated due to inefficiency of re-isolating bacteria from woody tissue, but this does show that the bottleneck effect is quite severe. Similar tight bottlenecks occur in other disease models such as *Vibrio cholerae* and *Listeria monocytogenes* infections in animal models (35–37). This does not negate the use of Tnseq to identify virulence genes, but can increase the rate of false positives in “essential” gene identification, and can require experimental modification to increase effectiveness of the approach (35). Re-isolation of large bacterial populations representing the population within a whole plant can also be challenging because the host plants are large and woody, and there is a limit to the plant tissue that can be used for isolation. It is possible that these practical limitations are the reason for high variability between replicate plants and a relatively short list of predicted “essential” genes when including only the genes predicted by the Bio-TRADIS pipeline that were found in all three replicates of a treatment group.

Overall, the largest category of predicted essential genes was hypothetical proteins, suggesting that the Tnseq approach could be used to identify candidate virulence genes in *X. fastidiosa*. Based on the outcome of experimental validation for three of the predicted essential genes, it is clear that at least some of the predicted essential genes impact fitness or disease manifestation, but are not strictly essential for survival. However, this is still a useful outcome in that it offers a starting point for further research, and contributes to the understanding of fitness contributions of different pathways. There were also several hemagglutinin genes that were identified in the “essential” gene predictions. Hemagglutinin adhesins are important for biofilm maturation, and attenuate virulence in *X. fastidiosa* (38, 39). It is possible that the sampling methodology (petiole samples at a late stage of infection) collected a higher proportion of biofilm and attached cells where adhesion defects would contribute to lower prevalence of mutants. tRNA genes were also heavily represented among the predicted essential genes *in planta*. This is similar to other mutagenesis studies that characterize bacterial essential genes due to the fundamental role of tRNA in protein synthesis (40, 41). Additionally, in *Ralstonia solanacearum* two tRNA genes were predicted to be essential during stress imposed by plant immune responses (42). In general, the same categories of genes were predicted to be essentialin both host plants. Because only a low number of predicted essential genes were found overall, the gene categories that show up in grapevine but not almond or vice versa could be an artifact of the small sample sizes and experimental variability. It is also relevant that the *X. fastidiosa* M23 strain causes significant disease in both grapevines and almonds, and the cultivars used in this study are both highly susceptible. Comparison of essential gene predictions using resistant or tolerant plant material is probably necessary to understand essential functions related to plant host range.

Overall, the Tnseq approach shows some promise for research of *X. fastidiosa* infection in plants but comes with some distinct limitations. A useful outcome of this study is that it shows how essential gene predictions performed using standard methods will capture genes with fitness and/or virulence defects, as well as those that are strictly essential for survival in this experimental scenario. Although identifying genes with fitness and/or virulence defects may be useful in its own right, if the goal is only to obtain genes that are truly essential, some experimental modifications would be necessary. The following are some suggestions for future research in this area based on this initial work.

### Recommendations for future *Xylella fastidiosa* Tnseq studies

1. **Developing more transposon mutagenesis options.** Commercially available Tn5 transposome complexes can be transformed efficiently into many strains of *X. fastidiosa* and have been use in research for many years (16). However, it would be beneficial to expand the available options for random transposon mutagenesis in this species for several reasons. First, commercial Tn5 transposome complexes are expensive, not necessarily universally available, and there is no guarantee that a commercial product will be around indefinitely. Additionally, Tn5 mutagenesis does not work as effectively in all strains of *X. fastidiosa* which limits research in certain *X. fastidiosa*-host plant combinations. Continued adaptation of other plasmid based random transposon mutagenesis protocols for *X. fastidiosa* transformation will facilitate a wider range of research, as well as improve longevity of the methods (60). An added benefit would be the ability to use a wider range of computational approaches for Tnseq data analysis, as many of these methods are developed for specific transposons such as *mariner* derivatives or mutant libraries with random barcoding (61, 62).
2. **Dealing with bottlenecks.** The bottleneck caused by *X. fastidiosa* inoculation into xylem tissue is quite severe, with <1% inoculated cells estimated to establish in the plant. This occurs in many other disease models as well, and there are a number of experimental design modifications that can mitigate this issue to some extent (35, 63). One option would be to inoculate sub-pools of transposon libraries into multiple different plants, perhaps with higher inoculum doses. This would increase initial sequencing costs but could reduce false positive predictions. Another option would be to create and test less diverse, defined mutant libraries. This approach would lose some of the ability to identify completely unknown virulence genes, but could still accelerate mutant testing compared with testing individual knockout mutants alone. Additional modifications such as different computational approaches have also been developed to tackle this issue (35, 63).
3. **Evaluating mutant competition.** Competition is an important aspect of Tnseq experiments since fitness of mutants competing in a pool of other mutants can be different than the fitness of a mutant inoculated individually. Testing competition requires creating mutants tagged with different selective markers which is possible in *X. fastidiosa*, but does require additional investment of time and resources, and would increase the size of plant inoculation experiments significantly. In some cases, fitness differences in Tnseq screening are consistent with direct competition assays, in other cases results are different depending on the role of the mutation and the screening conditions (64, 65). Regardless, competition assays can add important information to initial screening predictions produced by Tnseq.
4. **Targeted sampling.** *X. fastidiosa* is not uniformly distributed in infected plants which makes isolation of a bacterial population representative of the entire infection challenging. In future experiments it might be useful to focus on specific sections or types of plant tissue to answer more specific questions about bacterial colonization. For example, focusing bacterial isolation on stem tissues a certain distance from the point of inoculation to evaluate which mutants are best able to move systemically.

Tnseq methodology has been widely used in bacterial research over the last 15 years. However, only a small minority of these studies have been performed in plant hosts (63). Bacterial species such as *X. fastidiosa* with very specific lifestyles present some unique challenges for this approach, but at the same time present opportunities to refine and adapt screening methods such as Tnseq for a wider range of research. The goal of this study was to provide some initial insight into how this approach can work for *X. fastidiosa* research in plants, and we hope that others will contribute to building on this in the future.

## Data Availability Statement

Sequencing data from Tnseq experiments are deposited in the Sequence Read Archive under BioProject #PRJNA1111135. Bacterial strains, plasmids, mutants, and transposon libraries can be obtained by contacting the corresponding author.

## Acknowledgments

Additional support from United States Department of Agriculture, Agricultural Research Service appropriated project #2034-22000-014-000-D. Mention of trade names or commercial products in this publication is solely for the purpose of providing specific information and does not imply recommendation or endorsement by the U.S. Department of Agriculture. USDA is an equal opportunity provider and employer.

