## Supplemental File 1 for "In planta transposon sequencing for virulence gene identification in *Xylella fastidiosa*"

**Table S1. Bottleneck estimation**

| **Inoculum Concentration** | **Total CFU inoculated (Mean ± SD)** | **Total CFU recovered (Mean ± SD)** | **Bottleneck estimate**  **(Percent)** |
| --- | --- | --- | --- |
| 0.25 | 1.6E8 ± 2.4E7 | 7.8E+03 ± 6.2E+03 | 0.0047 ± 0.0037 |
| 5.0 | 7.2E12 ± 6.5E11 | 5.3E+05 ± 2.8E+05 | 7.3E-6 ± 3.E-6 |

**Table S2. Sequencing library yield and quality**

| **Library** | **Index** | **Yield (Mb)** | **Q30*** | **Mean Quality** |
| --- | --- | --- | --- | --- |
| PD3_1 | AGTCAA | 5,352 | 96.11 | 36.34 |
| PD3_2 | AGTTCC | 6,993 | 95.96 | 36.31 |
| PD3_3 | ATGTCA | 5,317 | 95.94 | 36.30 |
| GR_1 | CCGTCC | 5,659 | 96.08 | 36.34 |
| GR_2 | GTAGAG | 4,497 | 96.52 | 36.42 |
| GR_3 | GTCCGC | 6,479 | 96.12 | 36.34 |
| AL_1 | GTGAAA | 6,288 | 96.02 | 36.32 |
| AL_2 | GTGGCC | 5,455 | 95.99 | 36.31 |
| AL_3 | GTTTCG | 5,761 | 96.21 | 36.35 |

*% of bases (PF) with a quality score greater or equal to 30

**Figure S1. Sequence of Tn5 transposon**

>Tn5 Kan-2

CTGTCTCTTATACACATCTCAACCATCATCGATGAATTGTGTCTCAAAAT

CTCTGATGTTACATTGCACAAGATAAAAATATATCATCATGAACAATAAA

ACTGTCTGCTTACATAAACAGTAATACAAGGGGTGTTATGAGCCATATTC

AACGGGAAACGTCTTGCTCGAGGCCGCGATTAAATTCCAACATGGATGCT

GATTTATATGGGTATAAATGGGCTCGCGATAATGTCGGGCAATCAGGTGC

GACAATCTATCGATTGTATGGGAAGCCCGATGCGCCAGAGTTGTTTCTGA

AACATGGCAAAGGTAGCGTTGCCAATGATGTTACAGATGAGATGGTCAGA

CTAAACTGGCTGACGGAATTTATGCCTCTTCCGACCATCAAGCATTTTAT

CCGTACTCCTGATGATGCATGGTTACTCACCACTGCGATCCCCGGAAAAA

CAGCATTCCAGGTATTAGAAGAATATCCTGATTCAGGTGAAAATATTGTT

GATGCGCTGGCAGTGTTCCTGCGCCGGTTGCATTCGATTCCTGTTTGTAA

TTGTCCTTTTAACAGCGATCGCGTATTTCGTCTCGCTCAGGCGCAATCAC

GAATGAATAACGGTTTGGTTGATGCGAGTGATTTTGATGACGAGCGTAAT

GGCTGGCCTGTTGAACAAGTCTGGAAAGAAATGCATAAACTTTTGCCATT

CTCACCGGATTCAGTCGTCACTCATGGTGATTTCTCACTTGATAACCTTA

TTTTTGACGAGGGGAAATTAATAGGTTGTATTGATGTTGGACGAGTCGGA

ATCGCAGACCGATACCAGGATCTTGCCATCCTATGGAACTGCCTCGGTGA

GTTTTCTCCTTCATTACAGAAACGGCTTTTTCAAAAATATGGTATTGATA

ATCCTGATATGAATAAATTGCAGTTTCATTTGATGCTCGATGAGTTTTTC

TAATCAGAATTGGTTAATTGGTTGTAACACTGGCAGAGCATTACGCTGAC

TTGACGGGACGGCGGCTTTGTTGAATAAATCGAACTTTTGCTGAGTTGAA

GGATCAGATCACGCATCTTCCCGACAACGCAGACCGTTCCGTGGCAAAGC

AAAAGTTCAAAATCACCAACTGGTCCACCTACAACAAAGCTCTCATCAAC

CGTGGCGGGGATCCTCTAGAGTCGACCTGCAGGCATGCAAGCTTCAGGGT

TGAGATGTGTATAAGAGACAG

**Supplemental Figure S2. PCoA plots of predicted essential genes by treatment.**

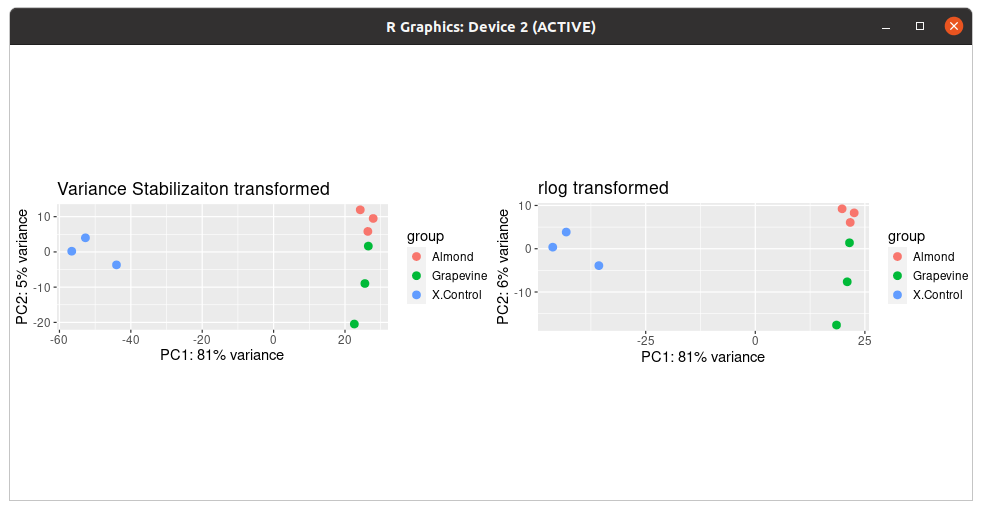

**Table S3. Bacterial strains and plasmids**

|  | Description | Source |
| --- | --- | --- |
| *Xylella fastidiosa* |  |  |
| M23 | Wild type *X. fastidiosa* subsp. *fastidiosa* | (1) |
| M23:Tn5 | Randon transposon mutant library, Kan^R^ | This study |
| ∆0359 | Targeted deletion mutant in XfasM23_0359, Kan^R^ | This study |
| ∆0360 | Targeted deletion mutant in XfasM23_0360, Kan^R^ | This study |
| ∆0972 | Targeted deletion mutant in XfasM23_0972, Kan^R^ | This study |
| ∆0359/0359+ | Complemented strain for ∆0359, Kan^R^, Gm^R^ | This study |
| ∆0360/0360+ | Complemented strain for ∆0360, Kan^R^, Gm^R^ | This study |
| ∆0972/0972+ | Complemented strain for ∆0972, Kan^R^, Gm^R^ | This study |
| *Plasmids* |  |  |
| pCR8/GW/TOPO | Cloning vector, Sp^R^ | ThermoFisher |
| pCR8-0359-kan | Mutagenesis construct for XfasM23_0359 | This study |
| pCR8-0360-kan | Mutagenesis construct for XfasM23_0360 | This study |
| pCR8-0972-kan | Mutagenesis construct for XfasM23_0972 | This study |
| pAX1/GW | Complementation vector for genomic insertion, Gateway® cloning compatible, Gm^R^ | (2, 3) |
| pAX1-0359 | Complementation construct for XfasM23_0359 | This study |
| pAX1-0360 | Complementation construct for XfasM23_0360 | This study |
| pAX1-0972 | Complementation construct for XfasM23_0972 | This study |
| pCRBlunt | Blunt cloning vector, ccdB counterselection, Kan^R^ | ThermoFisher |
| pCRBlunt-0360 | pCRBlunt containing XfasM23_0360, Kan^R^ | This study |
| pCRBlunt-0361 | pCRBlunt containing XfasM23_0361, Kan^R^ | This study |
| pCRBlunt-0360-0361 | pCRBlunt containing XfasM23_0360 and XfasM23_0361, Kan^R^ | This study |

**Table S4. Primer sequences**

| **Primer** | **Sequence** | **Description** | **Source** |
| --- | --- | --- | --- |
| 5GPWF | GGCTTTGACGACTGGTAGCT | Mutagenesis construct | This study |
| 5GPWRKAN | CAGCTGGCAATTCCGGGCTCAGCAGCTTGGGG | Mutagenesis construct | This study |
| 3GPWR | ATGTGTCTGTGGTGTAGGCG | Mutagenesis construct | This study |
| 3GPWFKAN | TCTTGACGAGTTCTTCTGAATGGACAACACTCTGACGGC | Mutagenesis construct | This study |
| KANF5GPW | CCCCAAGCTGCTGAGCCCGGAATTGCCAGCTG | Mutagenesis construct | This study |
| KANR3GPW | GCCGTCAGAGTGTTGTCCATTCAGAAGAACTCGTCAAGA | Mutagenesis construct | This study |
| 5PMSKF | AATCGCCGTCAGAGTGTTGT | Mutagenesis construct | This study |
| 5PMSKRKAN | AGCTGGCAATTCCGGCAGGCGGCCAGTATAC | Mutagenesis construct | This study |
| 3PMSKR | AACACCACCACCGAACAGTT | Mutagenesis construct | This study |
| 3PMSKFKAN | CTTGACGAGTTCTTCTGAGAAACACACGATGAAACGTATG | Mutagenesis construct | This study |
| KANF5PMSK | GTATACTGGCCGCCTGCCGGAATTGCCAGCT | Mutagenesis construct | This study |
| KANR3PMSK | CATACGTTTCATCGTGTGTTTCTCAGAAGAACTCGTCAAG | Mutagenesis construct | This study |
| 5M23-0972F | TAGGAATATCTAAACAAGCG | Mutagenesis construct | This study |
| 5M23-0972RKAN | CAGCTGGCAATTCCGGTTTTCCTAGTGTTTTCTTTACATA | Mutagenesis construct | This study |
| 3M23-0972R | TAGTTTGTATAAGTCTGCTG | Mutagenesis construct | This study |
| 3M23-0972FKAN | TCTTGACGAGTTCTTCTGATCTGAGAAATACATCAATAAA | Mutagenesis construct | This study |
| KANF5M23-0972 | TATGTAAAGAAAACACTAGGAAAACCGGAATTGCCAGCTG | Mutagenesis construct | This study |
| KANR3M23-0972 | TTTATTGATGTATTTCTCAGATCAGAAGAACTCGTCAAGA | Mutagenesis construct | This study |
| M23-0359-F | CCTTGCCAGCTTTCACCAAG | Mutant confirmation | This study |
| M23-0359-R | AATGGGATCGGGATGTCAGC | Mutant confirmation | This study |
| M23-0360-F | GGCGCTGACGTCAAGTACG | Mutant confirmation | This study |
| M23-0360-R | CTTCCGGCGTACCTTGGTAAT | Mutant confirmation | This study |
| M23-0972-F | ACACTCCTACCGTGCCCTAT | Mutant confirmation | This study |
| M23-0972-R | CGTCGTTTTGGTACGCGTTT | Mutant confirmation | This study |
| 0359-ORF-F | TTGTTGAATGCCTTTGTGGCG | Gene cloning for complementation | This study |
| 0359-ORF-R | TGACGGCTTTGGCATCTTCA | Gene cloning for complementation | This study |
| 0360-ORF-F | AGGTCCATTGTTAGGGCACG | Gene cloning for complementation | This study |
| 0360-ORF-R | AGCCGACCAATTCGCTGTAA | Gene cloning for complementation | This study |
| 0972-ORF-F | AGATGGGTACGAGAACTAGCA | Gene cloning for complementation | This study |
| 0972-ORF-R | TTTGATGGATTGTCAATAACGCA | Gene cloning for complementation | This study |
| Xf145-60F | TACATCGGAATCTACCTTATCGTG | qPCR quantification of *X. fastidiosa* |  |
| Xf145-60R | ATGCGGTATTTAGCGTAAGTTTC | qPCR quantification of *X. fastidiosa* |  |
| 0360-T-rev | ACGTTTCATCGTGTGTTTCCT | Expression in *E.coli* |  |
| 0361-AT-fwd | CGCCCCGAAGATATGAATGCT | Expression in *E.coli* |  |
| 0361-AT-rev | GTGCGACTAAACATGAAGGCG | Expression in *E.coli* |  |

**Table S5. Predicted essential genes for growth in different conditions**. List includes genes that are predicted to be essential in all three replicates of a specific condition but not in the other conditions.

| **Locus Tag** | **essential in** | **function** | **category** |
| --- | --- | --- | --- |
| XfasM23_0596 | Almond | cell division topological specificity factor MinE | cell division |
| XfasM23_1434 | Almond | lytic transglycosylase catalytic | conjugal transfer |
| XfasM23_1234 | Almond | putative conjugal transfer protein TrbL | conjugal transfer |
| XfasM23_1289 | Almond | ApaG domain-containing protein | efflux |
| XfasM23_1069 | Almond | CG31952 | hypothetical protein |
| XfasM23_0016 | Almond | hypothetical protein | hypothetical protein |
| XfasM23_0515 | Almond | hypothetical protein | hypothetical protein |
| XfasM23_0639 | Almond | hypothetical protein | hypothetical protein |
| XfasM23_0796 | Almond | hypothetical protein | hypothetical protein |
| XfasM23_0972 | Almond | hypothetical protein | hypothetical protein |
| XfasM23_1208 | Almond | hypothetical protein | hypothetical protein |
| XfasM23_1314 | Almond | hypothetical protein | hypothetical protein |
| XfasM23_1684 | Almond | hypothetical protein | hypothetical protein |
| XfasM23_1873 | Almond | hypothetical protein | hypothetical protein |
| XfasM23_1930 | Almond | hypothetical protein | hypothetical protein |
| XfasM23_1418 | Almond |  | non-coding |
| XfasM23_R0019 | Almond |  | non-coding |
| XfasM23_1109 | Almond | acetolactate synthase isozyme II small subunit | Amino acid biosynthesis |
| XfasM23_1119 | Almond | outer membrane lipoprotein Slp | outer membrane |
| XfasM23_1736 | Almond | ribosomal protein S21 | rRNA |
| XfasM23_0773 | Almond | general secretion pathway protein I | secretion |
| XfasM23_0360 | Almond | plasmid maintenance system killer | toxin-antitoxin system |
| XfasM23_R0015 | Almond | tRNA-Glu | tRNA |
| XfasM23_R0039 | Almond | tRNA-Gly | tRNA |
| XfasM23_R0048 | Almond | tRNA-Gly | tRNA |
| XfasM23_R0038 | Almond | tRNA-Ser | tRNA |
| XfasM23_R0040 | Almond | tRNA-Thr | tRNA |
| XfasM23_1044 | Grapevine | filamentous haemagglutinin outer membrane protein | hemagglutinin |
| XfasM23_2221 | Grapevine | filamentous haemagglutinin outer membrane protein | hemagglutinin |
| XfasM23_1331 | Grapevine |  | hemagglutinin |
| XfasM23_1018 | Grapevine | hypothetical protein | hypothetical protein |
| XfasM23_1022 | Grapevine | hypothetical protein | hypothetical protein |
| XfasM23_1043 | Grapevine | hypothetical protein | hypothetical protein |
| XfasM23_1210 | Grapevine | hypothetical protein | hypothetical protein |
| XfasM23_1211 | Grapevine | hypothetical protein | hypothetical protein |
| XfasM23_1326 | Grapevine | hypothetical protein | hypothetical protein |
| XfasM23_0994 | Grapevine | replication initiation factor | cell division |
| XfasM23_1209 | Grapevine | SNF2-related protein | phage |
| XfasM23_0370 | Grapevine | putative phage related protein | phage |
| XfasM23_R0009 | Grapevine |  | rRNA |
| XfasM23_R0012 | Grapevine |  | rRNA |
| XfasM23_1261 | Grapevine | addiction module antitoxin | toxin-antitoxin system |
| XfasM23_R0037 | Grapevine | tRNA-Arg | tRNA |
| XfasM23_1027 | PD3 | hypothetical protein | hypothetical protein |
| XfasM23_0368 | PD3 | hypothetical protein | hypothetical protein |
| XfasM23_0813 | PD3 | hypothetical protein | hypothetical protein |
| XfasM23_1047 | PD3 | hypothetical protein | hypothetical protein |
| XfasM23_1127 | PD3 | hypothetical protein | hypothetical protein |
| XfasM23_1142 | PD3 | hypothetical protein | hypothetical protein |
| XfasM23_R0008 | PD3 | tRNA-Ser | tRNA-Ser |
| XfasM23_R0017 | PD3 | tRNA-Phe | tRNA-Phe |
| XfasM23_R0023 | PD3 | tRNA-Ala | tRNA-Ala |

**Figure S3. *Xylella fastidiosa* quantification from almond plants (petiole samples) after 12 weeks post-inoculation**.

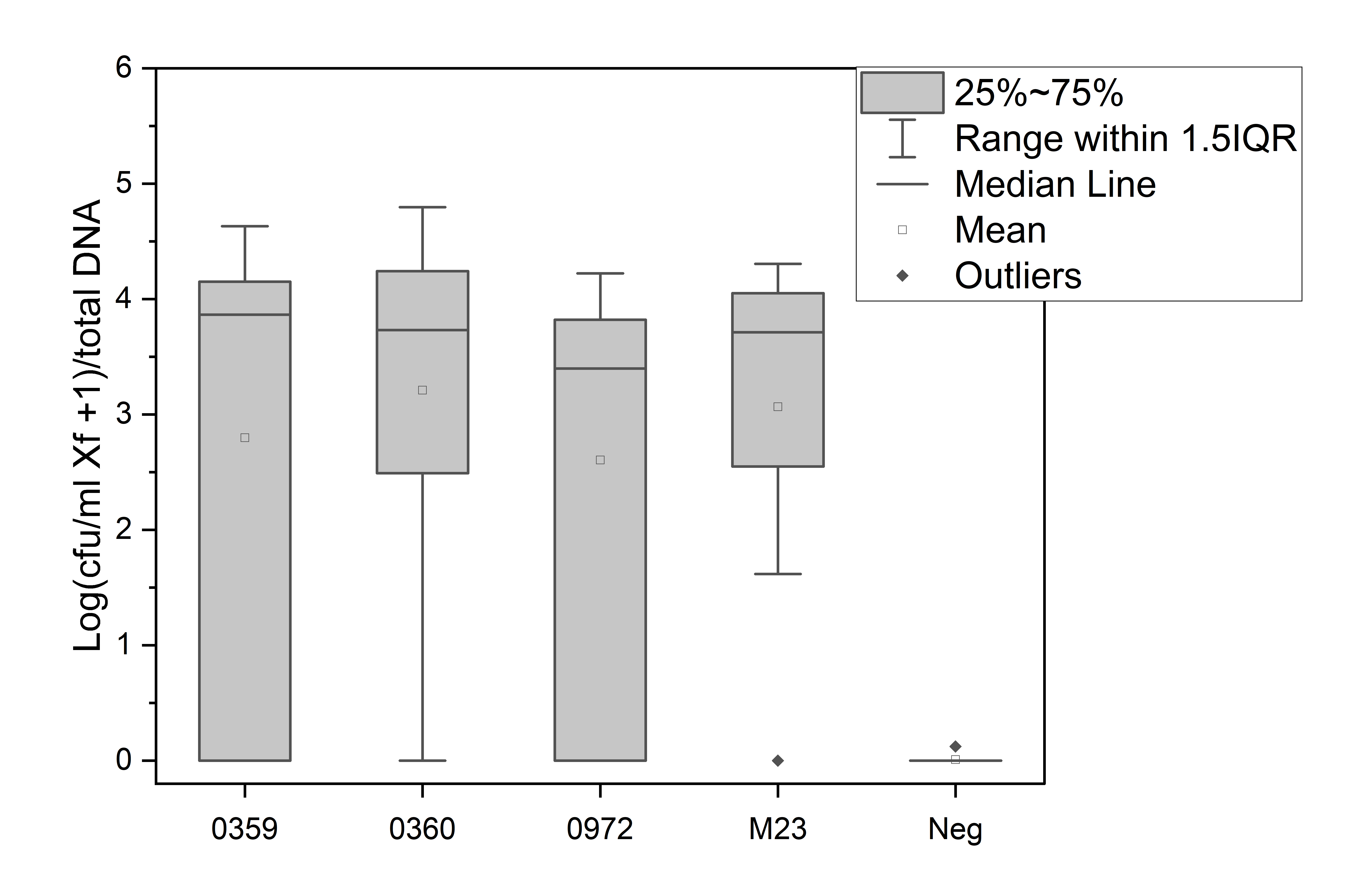
