## Supplemental File 3 for "In planta transposon sequencing for virulence gene identification in *Xylella fastidiosa*"

**Functional characterization of XfasM23_0972, XfasM23_0359, and XfasM23_0360**

**XfasM23_0972 and** **XfasM23_0359: Phage-associated proteins with unknown roles in disease.** Annotation for XfasM23_0359 is for a phage baseplate protein GPW/gp25 family protein, a gene commonly associated with prophage regions in *X. fastidiosa* and other bacterial species. XfasM23_0359 amino acid sequence shares 100% identity with a phage baseplate protein in other strains of *X. fastidiosa*, and 61.3% identity with a phage tail protein from *Escherichia coli* Phage vB_EcoM_ECOO78. Other similarity was with similar phage-associated proteins of the GpW/gp25 superfamily. Complete search results from the Uniprot database are shown in supplemental Figure S3.1. XfasM23_0972 is annotated as a hypothetical protein with very few functional or domain associations. Classification with InterPro identified a transmembrane domain but no other associated functions. Closest protein identities for XfasM23_0972 are 100% identity to hypothetical protein genes in other *X. fastidiosa* strains, and 66% identity to a phage-associated protein in *X. fastidiosa* Temecula1 strain. Other database matches were less than 50% identity. Complete Uniprot search results are available in Figure S3.2.

**XfasM23_0360: toxin component of a toxin-antitoxin system.** XfasM23_0360 is annotated as a HigB family toxin component of a Type II toxin-antitoxin system. This type of system typically consists of a protein toxin with endoribonuclease activity, and an adjacent protein antitoxin that directly interacts with the toxin to mitigate its toxic activity in the cell. The gene present directly downstream of XfasM23_0360 is the antitoxin component of the system (XfasM23_0361). To evaluate the role of these two genes together as a functional toxin-antitoxin system, XfasM23_0360 and XfasM23_0361 open reading frames were cloned into *E. coli* separately as well as together. The toxin (XfasM23_0360) expressed alone produced very few colonies (11 colonies total across 5 replicates), not significantly above what was found in the negative control (Fig. S3.3). Three of the toxin clones were sequenced with Sanger sequencing, and contained point mutations suggesting toxin activity may be reduced in the few clones that grew. The antitoxin (XfasM23_0361) alone produced the highest number of transformants, significantly more than both genes cloned together (Fig. S3.3). Sanger sequencing of three randomly selected clones of the antitoxin construct had no mutations compared with the reference sequences.

**Discussion**

From the list of predicted essential genes for *in planta* growth, three were characterized further using targeted deletions, plant inoculations, and complementation. All three of the deletion mutants tested showed decreased disease symptoms compared with the wild type. These genes were chosen for further analysis because they had no previously known function in *X. fastidiosa* virulence, and could provide additional insights into the disease process. XfasM23_0359, a phage-associated GpW/gp25 family protein has no known function in *X. fastidiosa*. Similar phage genes are associated with Type VI secretion systems in other bacterial species (1, 2), but no Type VI secretion genes have been found in the genus *Xylella* (3). More generally however, prophage insertions are associated with a large range of functions in plant pathogenic bacteria (4), including genome organization and rearrangement in *X. fastidiosa* (5, 6). Genome sequencing of several *X. fastidiosa* subsp. *multiplex* strains found high prevalence of phage components in multiple different *X. fastidiosa* sequence types (7). Often these prophages contain toxin-antitoxin systems as is the case with XfasM23_0359 and the XfasM23_0360/XfasM23_0361 system. Similar *X. fastidiosa* toxin-antitoxin systems have known functions in cell aggregation, growth *in planta*, and disease development (8–10). Others have no effect on disease, but are differentially expressed in response to temperature changes (11). In this case, it was the antitoxin component (XfasM23_0360) that was predicted as essential. Disruption of the antitoxin could reduce bacterial growth if toxin continued to be expressed, and this particular toxin could be more highly expressed in the plant environment than others. The hypothetical protein XfasM23_0972 is also potentially associated with a prophage insertion although the sequence-based evidence is not as definitive in this case. The gene directly adjacent (XfasM23_0971) is annotated as a zonular occludens toxin (Zot). Zot homologues are predicted pathogenicity factors in other strains of *X. fastidiosa* (12), and commonly associated with prophages in *Vibrio* and *Campylobacter* species (13, 14). Since many *X. fastidiosa* prophage associated genes are induced under stress conditions (5), there may be a general role of prophages in *X. fastidiosa* adaptation to changing environmental conditions, whether this is due to growth modulation by toxin-antitoxin systems or other regulatory pathways. In marine bacterial species, prophages are believed to facilitate long-term bacterial survival by slowing metabolism under nutrient restrictive conditions (15). In *X. fastidiosa*, there is evidence for several different strategies to reduce growth rate and attenuate disease in plants which could be facilitated by prophage sequences in a similar manner (10, 16, 17). Specific phages in *Ralstonia solanacearum* can alter phenotypes associated with virulence and survival such as twitching motility and cellular aggregation (18). Although no differences in aggregation was observed in the *X. fastidiosa* mutants in this study, there could also be contributions of *X. fastidiosa* prophage gene expression to these phenotypes *in planta*. Overall, the essential gene predictions highlighted some genes that contribute to disease and had not previously been characterized in *X. fastidiosa*. With additional improvements to the methodology, this approach could be useful to further identify virulence genes in different *X. fastidiosa*-plant host interaction scenarios.

**Figure S3.1. Uniprot search results for XfasM23_0359**

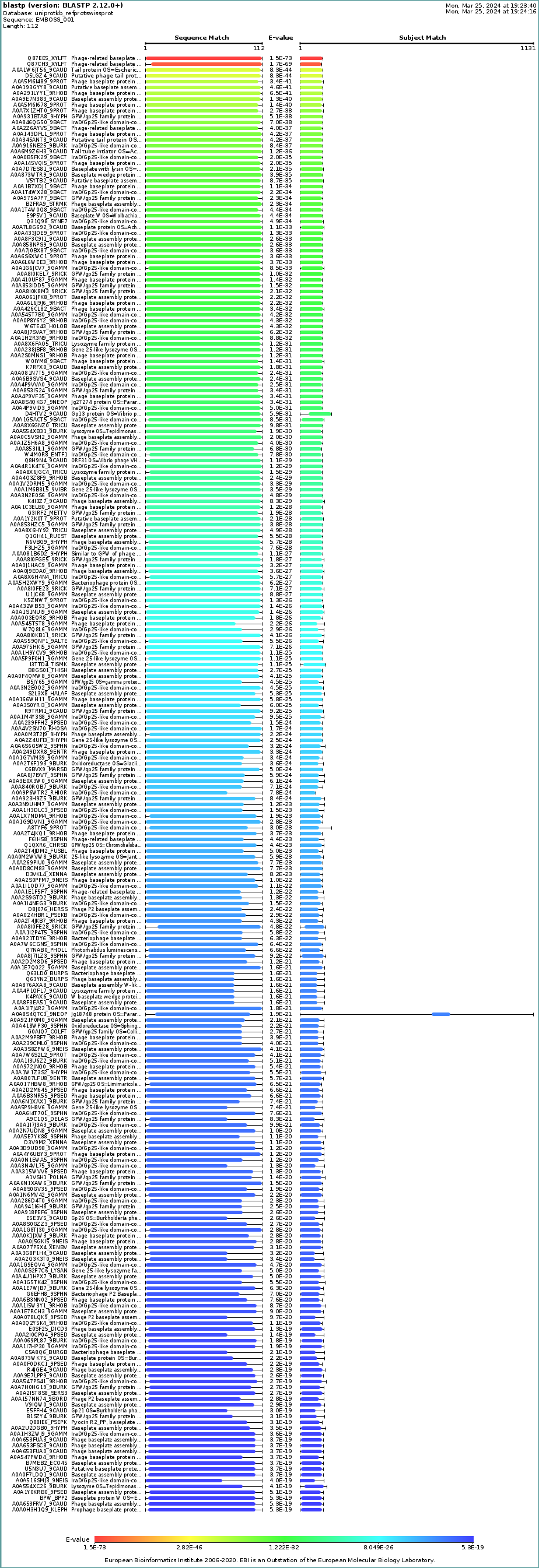

**Figure S3.2. Uniprot search results for XfasM23_0972**

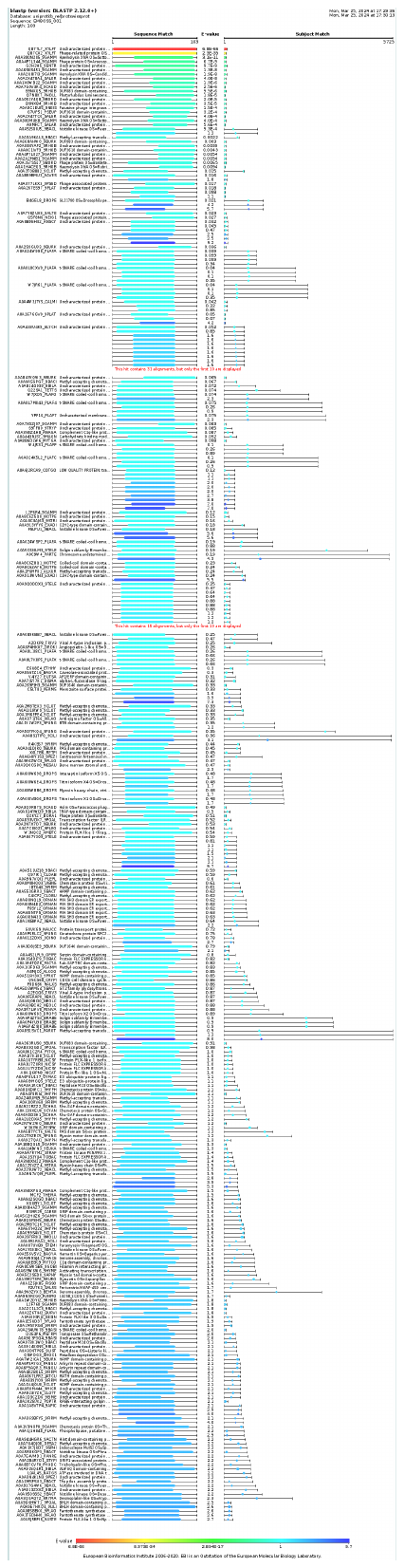

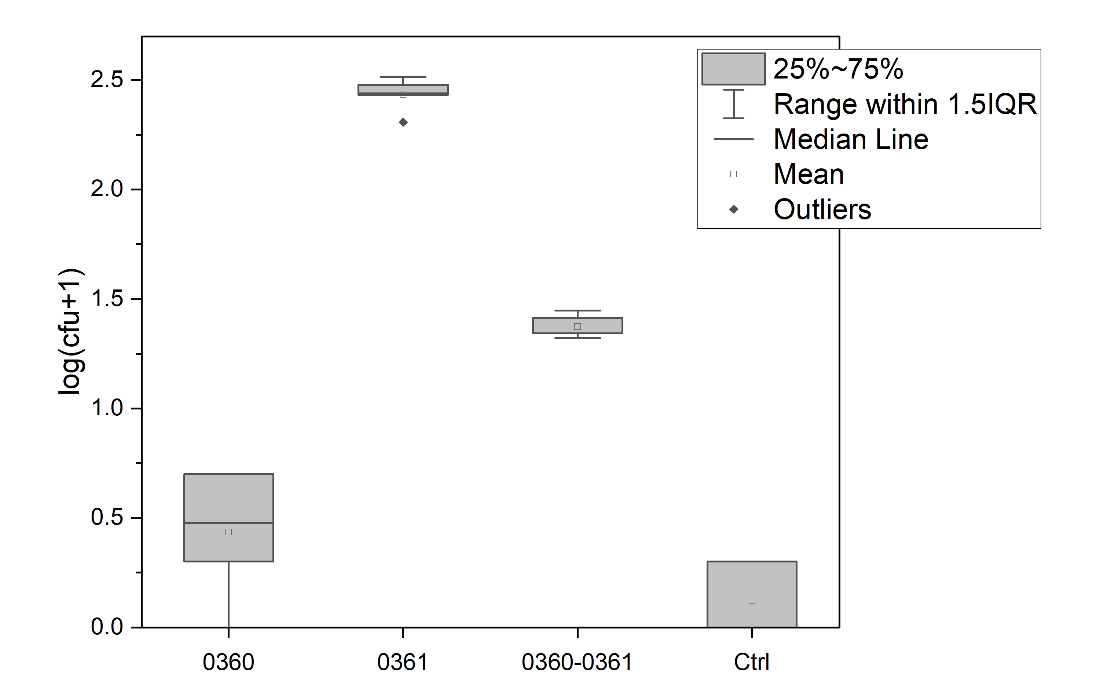

**Figure S3.3. Colonies from XfasM23_0360/XfasM23_0361 toxin-antitoxin gene cloning in *E. coli*.** Number of transformations was quantified after cloning open reading frames of XfasM23_0360 (toxin), XfasM23_0361 (antitoxin), and the whole toxin-antitoxin system (XfasM23_0360/XfasM23_0361) into plasmid pCRBlunt in *E. coli*. Five replicates of transformations were performed. Negative control (Ctrl) consisted of cloning reactions with no DNA insert.
